# Analogs of TIQ-A as inhibitors of human mono-ADP-ribosylating PARPs

**DOI:** 10.1101/2021.08.30.458193

**Authors:** Mirko M. Maksimainen, Sudarshan Murthy, Sven T. Sowa, Albert Galera-Prat, Elena Rolina, Juha P. Heiskanen, Lari Lehtiö

## Abstract

The scaffold of TIQ-A, a previously known inhibitor of human poly-ADP-ribosyltransferase PARP1, was utilized to develop inhibitors against human mono-ADP-ribosyltransferases through structure-guided design and activity profiling. By supplementing the TIQ-A scaffold with small structural changes, based on a PARP10 inhibitor OUL35, selectivity changed from poly-ADP-ribosyltransferases towards mono-ADP-ribosyltransferases. Binding modes of analogs were experimentally verified by determining complex crystal structures with mono-ADP-ribosyltransferase PARP15 and with poly-ADP-ribosyltransferase TNKS2. The best analogs of the study achieved 10 – 20-fold selectivity towards mono-ADP-ribosyltransferases PARP10 and PARP15 while maintaining micromolar potencies. The work demonstrates a route to differentiate compound selectivity between mono- and poly-ribosyltransferases of the human ARTD family.

## 1. Introduction

Human PARP enzymes of the diphtheria toxin-like ADP-ribosyltransferasefamily (ARTD)^1^, form a group of 17 enzymes sharing a catalytic domain carrying out the post-translational protein modification called ADP-ribosylation.^2,3^ Based on the nature of the resulting modification, the family members are classified as poly-ADP-ribosyltransferases (poly-ART) or mono-ADP-ribosyltransferases (mono-ART). Poly-ARTs PARP1-2, TNKS1-2 produce linear and branched polymers of ADP-ribose (PAR) to target proteins, whereas most of the enzymes in the family are actually mono-ARTs which transfer single ADP-ribose units to a target. ADP-ribosylation regulates vital cellular processes like transcription, signaling and DNA repair and thus PARPs are possible therapeutic targets against human diseases, especially cancer.

The catalytic PARP domain contains a substrate NAD^+^ binding site comprising both nicotinamide and adenosine binding pockets. Due to that, PARP inhibitors have been developed to target either one of the pockets or both. To date, clear majority of known PARP inhibitors are nicotinamide mimetics and the inhibitor development is more advanced against poly-ART members than mono-ART as the best PARP1-2 inhibitors olaparib, rucaparib, niraparib and talazoparib are already in clinical use^4,5^. In addition, tankyrase inhibitor development has recently resulted in lead compounds showing promising antitumor activities^6,7^. In the case of mono-ADP-ribosyltransferases, approximately half of the members are lacking selective inhibitors. Based on the available literature, selective inhibitors have been discovered for PARP7^8^, PARP10^9^, PARP11^10^, PARP14^11^and PARP15^12^ but their usability in clinical treatments remains to be elucidated. A generally known challenge in the PARP inhibitor development is gaining of selectivity due to the conserved catalytic domain of PARPs.

Thieno[2,3-c]isoquinolin-5(4H)-one, also known as TIQ-A (**1**) was originally developed as a potential anti-ischemic agent inhibiting PARP1^13,14^. We recognized, while evaluating inhibitors against tankyrases, that **1** is indeed also a potent TNKS inhibitor^15^. Notably, many early PARP inhibitors, including approved drugs, suffer from a lack of selectivity between the PARP family members^16^. In the case of **1**, its poor selectivity is obvious as we have previously shown the compound is also efficient to inhibit mono-ART PARP15^9^. **1** belongs to nicotinamide mimicking inhibitors and partially resembles OUL35, a selective inhibitor of PARP10^17^. From the structure activity relationship study of PARP10 inhibitors^18^ we identified an analog (**2**), which together with **1** formed a basis for our attempts to optimize the scaffold **1** in such a way that it would have selectivity towards different PARPs. A key feature of **2** is that it extends towards the acceptor site of the enzyme where protein to be modified is expected to bind. This region is more polar in poly-ARTs containing an active site glutamate required for PAR elongation reaction, while the corresponding residue in most mono-ARTs is a hydrophobic one. By using a selected set of both poly- and mono-ART PARPs as examples we demonstrate that the selectivity profile of **1** can be changed through appropriate substitutions and the differences can be explained to some extent using complex crystal structures of the analogs and a poly-ART (TNKS2) or a mono-ART (PARP15).

## 2. Materials and Methods

### 2.1. Chemistry

The compounds **3** – **8** were synthesized during our previous studies.^19^ The syntheses of new molecules **9** and **10** are described below. All commercial starting materials and reagents were used without purification. The solvents were dried with appropriate molecular sieves when needed. The reaction progress was monitored with silica gel-coated aluminum TLC sheets. The chemical structures of **9** and **10** were characterized using ^1^H NMR, ^13^C NMR, and HRMS measurements. ^1^H and ^13^C NMR assignments of **9** and **10** are presented in the Figures S1-S6. Purities of **4** – **8** were assessed using LCMS by UV absorbance and are ≥95% unless otherwise stated.

#### Methyl 3-(3-ethoxyphenyl)thiophene-2-carboxylate (9)

Compound **3** (137.2 mg; 586 mmol) and K_2_CO_3_ (292.0 mg; 2.11 mmol) were placed in a reaction tube. The sealed tube was evacuated and backfilled with argon three times. The mixture was stirred and heated in an oil bath (65°C). Diethyl sulfate (0.12 mL; 918 mmol) was added. The reaction was allowed to proceed 18 h. Water (10 mL) and EtOAc (10 mL) were added. The aqueous phase was extracted with EtOAc (2 x 10 mL) and the combined organic phase was filtered through a thin pad of silica, which was rinsed with EtOAc. After evaporation, the sticky residue was washed with water (3 x 10 mL) and dried under vacuum. The procedure afforded **9** as a sticky solid (142.0 mg) in 92% yield. ^1^H NMR (400 MHz, CDCl_3_) *δ* ppm 1.45 (t, J=7.0 Hz, 3H), 3.80 (s, 3H), 4.08 (q, J=7.0 Hz, 2H), 6.94 (ddd, J=8.3, 2.5, 0.9 Hz, 1H), 7.03– 7.07 (m, 2H), 7.10 (d, J=5.1 Hz, 1H), 7.31–7.35 (m, 1H), 7.50 (d, J=5.1 Hz, 1H). ^13^C NMR (100 MHz, CDCl_3_) *δ* ppm 14.8, 51.8, 63.3, 114.0, 115.3, 121.5, 126.9, 128.7, 130.1, 131.5, 136.8, 148.3, 158.3, 162.3. HRMS (ESI) m/z: [M + H]^+^ Calcd for C_14_H_15_O_3_S 263.0736; Found 263.0737.

#### 8-Ethoxythieno[2,3-c]isoquinolin-5(4H)-one (10)

Compound **9** (142.0 mg, 541 mmol) was added to a 100 mL round-bottom flask. EtOH (6.8 mL), H_2_O (6.8 mL), and ground NaOH (433.0 mg, 10.8 mmol) were added and the reaction mixture was refluxed for 90 min. CH_2_Cl_2_ (15 mL) was added to the cooled mixture. Aq. HCl (37%) was added dropwise until the pH value reached 2. The aqueous phase was extracted with CH_2_Cl_2_ (2 x 15 mL). The combined organic phase was washed with water (2 x 10 mL), dried (Na_2_SO_4_) and filtered. The solvent was evaporated under vacuum to give the intermediate carboxylic acid as a white powder (136.0 mg; >99%). The acid was transferred into a 50 mL two-neck round-bottom flask and dry toluene (3.75 mL) was added. The mixture was stirred at 80°C under argon. Thionyl chloride (0.07 mL, 960 mmol) and dry DMF (few drops) were added. After 105 min, the solvent and excess reagent were evaporated. THF (3.0 mL) was added under argon and the mixture was stirred at 0°C. NaN_3_ (52.7 mg, 811 mmol) in H_2_O (0.45 mL) was added slowly to the mixture. After 20 min, ice-water (10 mL) and CH_2_Cl_2_ (10 mL) were added. The phases were separated and the aqueous layer was extracted with CH_2_Cl_2_ (2 x 10 mL). The combined organic phase was dried (Na_2_SO_4_) and filtered. The evaporation residue was dissolved in 1,2-dichlorobenzene (4.5 mL) and added to hot (210°C) 1,2-dichlorobenzene (6 mL) in a 50 ml two-neck round-bottom flask equipped with a reflux condenser and an argon balloon. The reaction mixture was stirred at 210°C for 20 h. The evaporation residue was dissolved in a mixture of toluene and EtOAc (1:1) and subjected to a flash chromatography using EtOAc as an eluent. The collected solid was washed with n-hexane (3 × 5 mL), and dried to afford **10** as light-grey powder (104.5 mg) in 79% yield. ^1^H NMR (400 MHz, CDCl_3_) δ ppm 1.51 (t, J=7.0 Hz, 3H), 4.22 (q, J=7.0 Hz, 2H), 7.00 (d, J=5.6 Hz, 1H), 7.08 (dd, J=8.9, 2.3 Hz, 1H), 7.26–7.27 (m, 1H), 7.43 (d, J=5.6 Hz, 1H), 8.44 (d, J=8.9 Hz, 1H), 11.37 (br s, 1H). 13C NMR (100 MHz, (CD_3_)_2_SO) δ ppm 14.5, 63.7, 105.6, 115.0, 116.2, 116.9, 117.6, 121.5, 129.9, 135.6, 141.3, 161.0, 162.2. HRMS (ESI) m/z: [M + H]^+^ Calcd for C_13_H_11_NO_2_S 246.0583; Found 246.0583.

### 2.2. Protein expression and purification

All the proteins used in this study were expressed in *E. coli* and purified using our previously reported protocols.^17^ TNKS2 constructs were cloned into pNIC-MBP expression vectors and an additional purification on an MBPTrap HP 5 ml column (GE Healthcare) prior to cleavage with TEV protease was performed.^20^ Details of the constructs for each PARP enzyme are listed in Table S1.

### 2.3. Activity assay

Dose response experiments were performed using our previously reported activity assay for PARP enzymes.^9,21^ Half-log dilutions for the compounds were used and reactions were carried out in quadruplicates. IC_50_ curves were fitted using sigmoidal dose response curve (four variables) in GraphPad Prism version 5.04 (GraphPad Software). Details of the assay conditions of different PARP enzymes are available in Table S1.

### 2.4. Crystallization

Crystallization of TNKS2 and soaking with **7, 8** and **10** was done as previously described.^22^ PARP15 was co-crystallized with **1, 2, 4, 5, 6, 7, 8** and **10** utilizing the existing crystallization conditions for PARP15.^23^ Compounds were dissolved in DMSO and 1.5 μl of resulting 10 mM solution was gently mixed with 20 μl of 10.5 mg/ml PARP15 and incubated for 30 s at 20°C for crystallization. 100 - 150 nl of a protein-compound solution was mixed well with 75 – 100 nl of solution consisting of 0.2 M NH_4_Cl pH 7.5, 16 – 20% (w/v) PEG 3350 using the Mosquito crystallization robot (SPT Labtech) and 3 well low profile UVP (Swissci) for sitting drop vapour diffusion. Crystallization plates of PARP15 and TNKS2 were incubated at +20°C and +4°C, respectively. All plates were imaged using the RI54 imager (Formulatrix) at the respective temperature and monitoring of crystallization was performed using the IceBear software.^24^ PARP15 crystals were obtained in 24 h while TNKS2 crystals were obtained in 5 days.

### 2.5. Data collection, processing and refinement

Prior to data collection, PARP15 crystals were cryoprotected with a solution consisting of 0.2 M NH_4_Cl pH 7.5, 30% (v/v) MPD (2-methyl-2,4-pentanediol). TNKS2 crystals were soaked in reservoir solution containing 20% glycerol. The X-ray diffraction data were collected at the synchrotron facilities of DLS (Didcot, UK) and ESRF (Grenoble, France). All data were processed and scaled with the XDS program.^25^ The PARP15 and TNKS2 complex structures were solved by molecular replacement with Molrep^26^ (from the CCP4 package^27^) using the previously reported structures of PARP15 (PDB accession code 3BLJ^23^) and TNKS2 (PDB accession code 5OWS^28^).The PARP15 and TNKS2 models were refined with Refmac5^29^ (from the CCP4 package^27^). Visualization and building of all models were performed using Coot.^30^ The residues in PARP15 and TNKS2 models were numbered according to the canonical sequences of UniProt^31^ entries Q460N3-1 and Q9H2K2-1, respectively. Data collection and refinement statistics are shown in the Table S2.

## 3. Results

### 3.1. Synthesis

Compound **3** was alkylated with diethyl sulfate using solvent-free Williamson synthesis^32^ which gave **9** in high yield (Scheme 1). 8-Ethoxythieno[2,3-c]isoquinolin-5(4H)-one **10** was synthesized following the method developed during our previous studies.^19^ The base catalyzed hydrolysis of the ester group of **9** resulted an acid that was converted to its acid chloride with thionyl chloride and catalytic amount of dry DMF. Reaction with NaN_3_ gave the corresponding carbonyl azide, which readily underwent Curtius rearrangement at elevated temperature in 1,2-dichlorobenzene, affording **10** in good yield.

**Scheme 1^a^.**
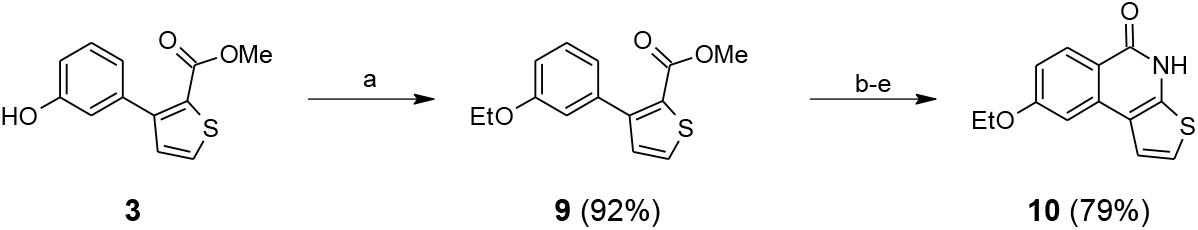
Synthesis of 8-ethoxythieno[2,3-c]isoquinolin-5(4H)-one (**10**) ^**a**^Reagents and conditions: (a) diethyl sulfate, K_2_CO_3_, 65°C, 18 h; (b) NaOH, EtOH, H_2_O, reflux, 90 min; (c) SOCl_2_, toluene, DMF, 80°C, 105 min; (d) NaN_3_, THF, H_2_O, ice bath, 20 min; (e) 1,2-dichlorobenzene, 210°C, 20 h.

### 3.2. Inhibitor design and structural studies

Compound **1** is a nicotinamide mimicking compound, which we have previously reported to inhibit human TNKS2, a poly-ART, at a nanomolar potency (IC_50_ = 24 nM).^15^ In addition, while developing an activity-based assay for human mono-ADP-ribosyl-transferases,^9^ we saw that the compound also inhibited PARP15 (IC_50_ = 232 nM) and modestly PARP10 (41% inhibition at 10 μM). The relatively high potency against PARP15 led us to hypothesize if **1** could be used as a scaffold to develop a selective inhibitor against mono-ADP-ribosylating PARP enzymes. However, based on the clear evidences on its unselectivity against human PARPs seen by us and also Wahlberg *et al*.^16^ (Table 1), the scaffold needed to be modified rationally to change the selectivity towards mono-ADP-ribosyltransferases. To facilitate the compound design, we determined the PARP15 crystal structure in complex with **1** (Fig. 1A) that showed the compound similarly bound into the nicotinamide binding pocket with three hydrogen bonds created by G560 and S599 compared to previously reported TNKS2 structure where three hydrogen bonds are created by G1032 and S1068 (Fig. 1B).^15^

**Figure 1:**
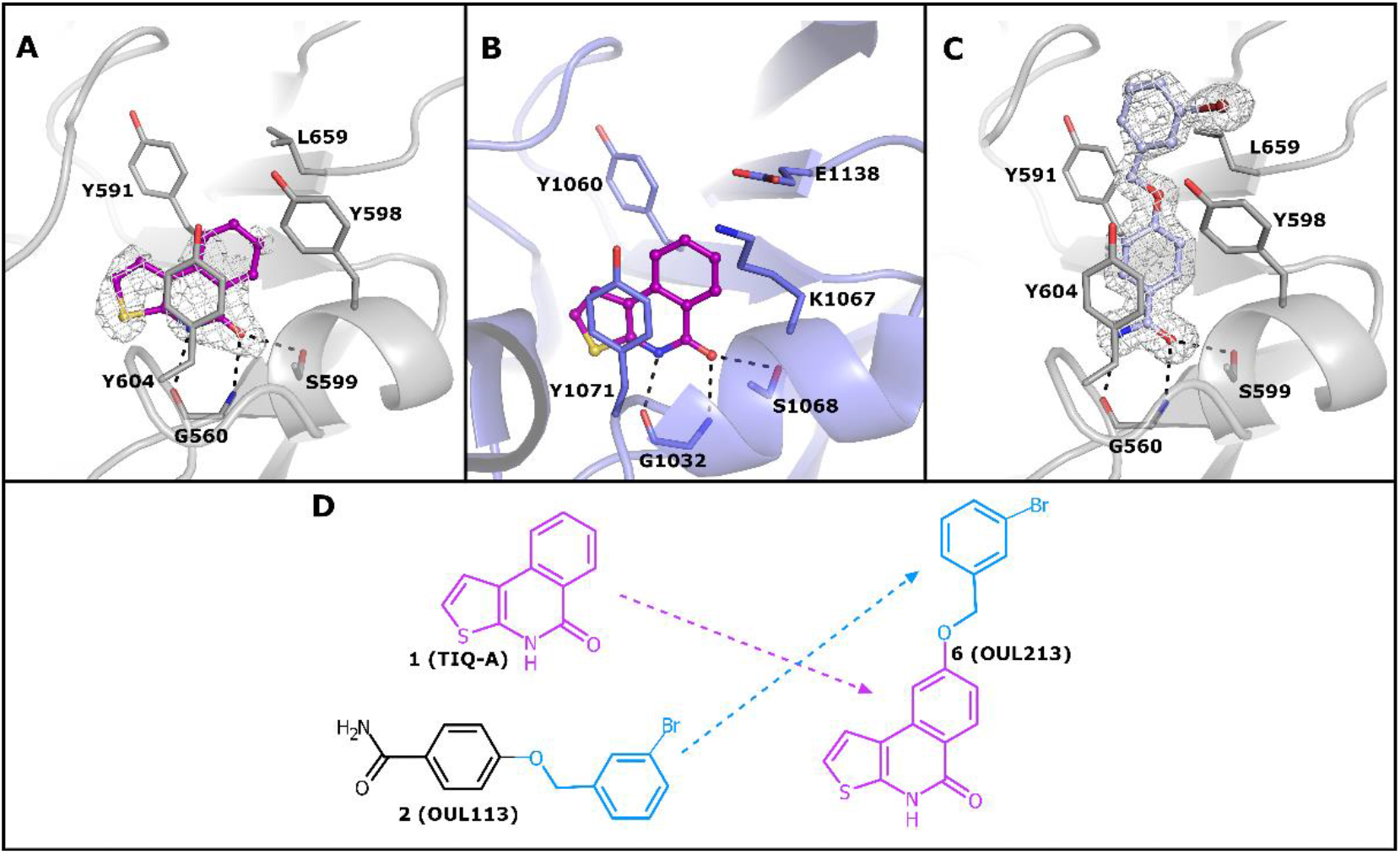
Inhibitor design based on the complex structures of PARP15 and TNKS2. **A)** PARP15 in complex with **1** (purple) **B)** TNKS2 in complex with **1** (PDB id 4AVW)^15^ **C)** PARP15 in complex with **2** (light blue) **D)** Design to merge **1** and **2** to create a chimera **6**. Sigma A weighted omit Fo-Fc electron density maps covering the ligands in A and C are contoured at 3.0 σ and colored in white. Ligands are presented as ball-stick models while the residues of PARP15 and TNKS2 are presented as stick models and colored in grey and blue, respectively. The secondary structures of PARP15 and TNKS2 are presented as a grey and blue cartoon models, respectively. The hydrogen bonds are shown as black dashed lines

**Table 1.** Profiling of the compounds against a panel of human ARTs

| ID | PARP2<br>IC<sub>50</sub><br>(pIC<sub>50</sub>±SEM) | TNKS2<br>IC<sub>50</sub><br>(pIC<sub>50</sub>±SEM) | PARP10<br>IC<sub>50</sub><br>(pIC<sub>50</sub>±SEM) | PARP14<br>IC<sub>50</sub><br>(pIC<sub>50</sub>±SEM) | PARP15<br>IC<sub>50</sub><br>(pIC<sub>50</sub>±SEM) | PDB codes of<br>crystals structures<br>(Enzyme) |
| --- | --- | --- | --- | --- | --- | --- |
| 1 | 210 nM* | 24 nM* | >10 μM* | > 10 μM | 230 nM* | 7OQQ (PARP15) |
| 2 | >100 μM | >100 μM | 2.4 μM | 630 nM | 1.2 μM | 7OSP (PARP15) |
| 4 | 87 μM | 57 μM | > 10 μM | > 100 μM | > 10 μM | 7OSS (PARP15) |
| 5 | 1.2 μM | 130 nM<br>(6.89±0.055) | 4.5 μM | > 100 μM | 1.4 μM | 7OSX (PARP15) |
| 6 | 60 μM | 20 μM | > 10 μM | > 100 μM | >> 10 μM | 7OTF (PARP15) |
| 7 | 19 μM | 22 μM | 2.4 μM | > 100 μM | 1.1 μM<br>(5.95±0.12) | 7OTH (PARP15)<br>7OLJ (TNKS2) |
| 8 | 1.8 μM | 160 nM<br>(6.86±0.08) | 780 nM | 52 μM | 527 nM<br>(6.28±0.03) | 7OUW (PARP15)<br>7OM1 (TNKS2) |
| 10 | >100 μM | 4.7 μM | 920 nM | > 10 μM | 2.1 μM | 7OUX (PARP15)<br>7OMC (TNKS2) |

\*Data from literature: PARP2<sup>16</sup>, TNKS2<sup>15</sup>, PARP10<sup>9</sup> and PARP15<sup>9</sup>

We thought that an extension of the scaffold towards the so-called acceptor site identified by Ruf *et al*.^33^, would change the selectivity towards mono-ADP-ribosyl-transferases as we have previously seen in the case of PARP10 inhibitors^17,18^. We chose **2** as a partner for **1** because the compound showed a reasonable potency against PARP15 (IC_50_ = 1.2 μM) and importantly did not show activity against poly-ARTs such as PARP2 and TNKS2 (Table 1). We determined the PARP15 crystal structure in complex with **2** that showed that the compound is bound with the same hydrogen bond interactions as **1** and it indeed extends towards the acceptor site (Fig. 1C). Especially regarding the inhibitor design, the bromophenyl moiety of **2** is clearly oriented towards the acceptor site. These observations encouraged us to merge the 3-bromophenylmethoxy from **2** to C-8 of the scaffold **1** (Fig. 1D).

In order to implement the idea, we started chemical synthesis^19^ from which we first obtained **4** and **5** before the desired compound **6**. The profiling of the compounds against PARP2, TNKS2, PARP10, PARP14, and PARP15 showed that **4** was not very potent against any enzyme whereas **5** inhibited PARP2, PARP10, and PARP15 with micromolar potencies, being most potent against TNKS2 (IC_50_ = 115 nM, Table 1). Like **4**, the desired compound **6** was not potent against any PARP. In fact, the compound had at least 100-fold and 20-fold less activity compared to **1** and **2**, respectively. Despite of low potencies of **4** and **6**, we were able to determine PARP15 complex crystal structures, which revealed that all three compounds bind to PARP15 as we had hypothesized (Fig. 2A-C). The superimposition of **1, 2** and **6** based on the PARP15 complex structures showed small differences between the positions of the compounds (Fig. 2D). The sulfur atoms of **1** and **6** had 0.7 Å distance between them while the bromine atoms of **2** and **6** had 0.8 Å distance and these subtle changes in the compound orientations could explain the poor potency of **6**.

**Figure 2:**
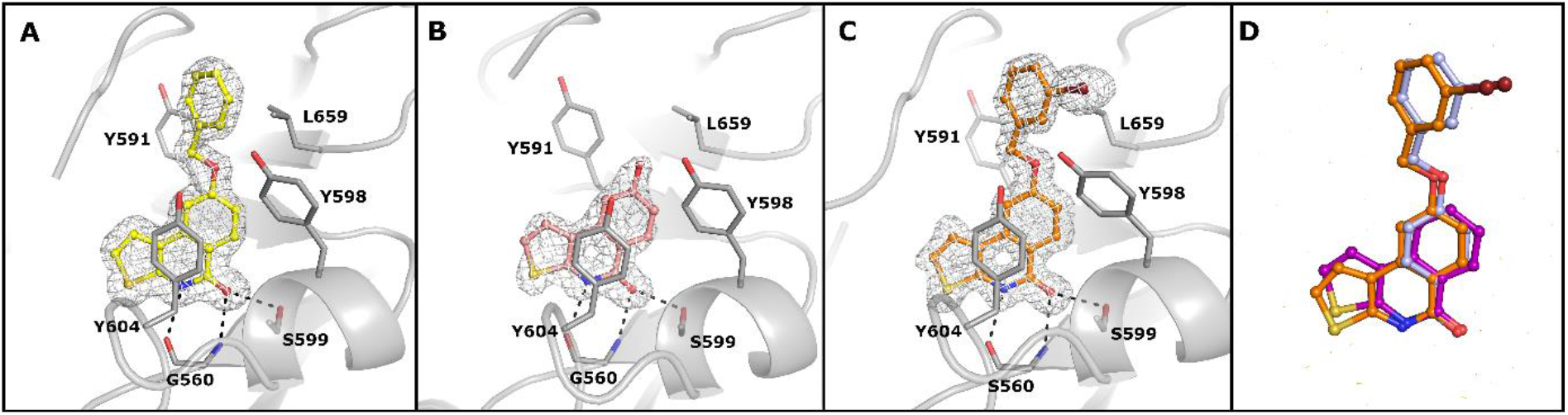
PARP15 crystal structures in complex with **A) 4** (yellow) **B) 5** (pink) **C) 6** (orange) **D)** Superimposition of **1** (purple), **2** (light blue) and **6** (orange) based on their PARP15 complex structures. For clarity, PARP15 is left out from the panel. The structures and electron density maps are presented as in Figure 1.

As the benzyloxy moieties in the scaffold **1** clearly hindered the inhibition, we rationalized that smaller substituents could work better. We tested analogs containing isopropoxy (**7**), methoxy (**8**), and ethoxy (**10**) substituents at C-8 of the scaffold **1. 7** showed approximately 10-fold selectivity for PARP10 (IC_50_ = 2.4 μM) and 20-fold selectivity for PARP15 (IC_50_ = 1.1 μM) over the poly-ARTs. **8** showed good potencies against PARP15 (IC_50_ = 527 nM) and PARP10 (IC_50_ = 780 nM) but lacked selectivity as it was also very active against TNKS2 (IC_50_ = 160 nM). **10** was not good as **7** and showed approximately 5-fold selectivity for PARP10 and 2-fold selectivity for PARP15 (Table 1). The complex crystal structures of **7, 8**, and **10** showed that binding of the compounds is highly similar in both PARP15 and TNKS2 (Fig. 3). The orientations of methoxy and ethoxy moieties in PARP15 are clearly defined by the electron density maps while the isopropoxy moiety of **7** lacks clear electron density indicating mobility of the group in the binding pocket (Fig. 3A-C). In TNKS2, the compounds are well defined by electron density maps (Fig. 3D-F). A comparison of TNKS2 crystal structures reveal significant conformational changes of the sidechain of the catalytic residue, E1138 (Fig. 1B, 3D-F). This small rotation of E1138 is caused by the hydrophobic substituents that explain lower potency of **7** and **10** for TNKS2 (Table 1).

**Figure 3:**
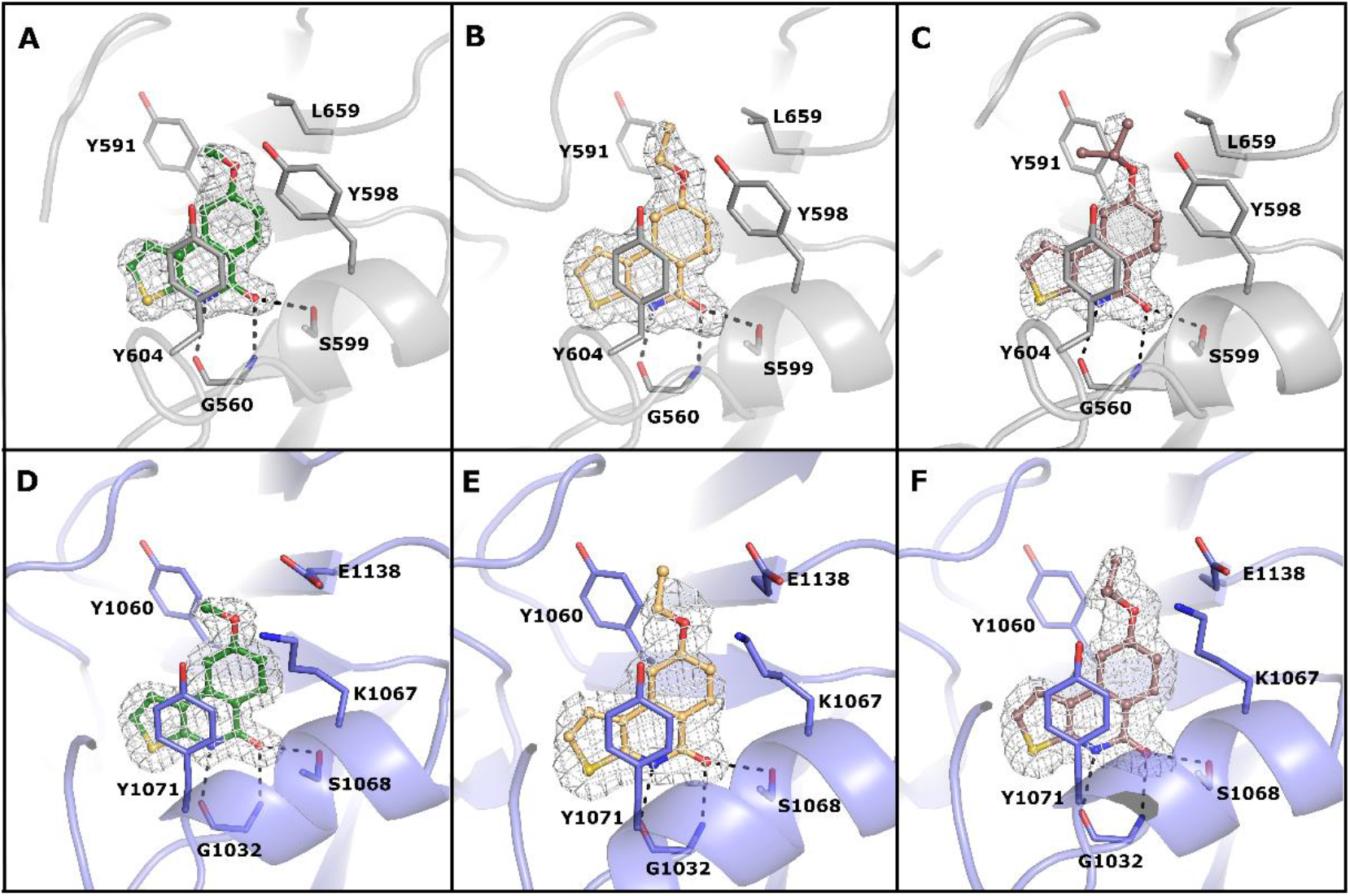
Comparison of PARP15 and TNKS2 complex crystal structures A) PARP15 in complex with **8** B) PARP15 in complex with **10** C) PARP15 in complex with **7** D) TNKS2 in complex with **8** E) TNKS2 in complex with **10** F) TNKS2 in complex with **7**. The ligands **8, 10** and **7** are colored in green, light orange and brown, respectively. Sigma A weighted omit Fo-Fc electron density maps covering the ligands are presented as in Fig. 1 and 2.

## 4. Discussion

Our aim was to develop selective inhibitors for human mono-ART class of PARP enzymes by utilizing the scaffolds of **1** and **2** and merging promising features through chemical synthesis. However, despite that the binding mode of hybrid analogs was experimentally verified to be same as planned, the direct combining the unique features of **1** and **2** did not work as relatively large benzyloxy moieties at C-8 of scaffold **1** resulted in significant loss of potency for all the PARPs used in the study. This could possibly be a result of subtle changes in the binding modes as in the co-crystal structures these analogs still maintained a highly similar binding mode (Fig. 2).

Instead, compounds **5, 7, 8** and **10** with smaller alkyloxy moieties at the same position showed micromolar or submicromolar potencies for both poly- and mono-ART enzymes. **7** containing an isopropoxy group showed the best selectivity (20-fold) for PARP15 in comparison to PARP2 and TNKS2 demonstrating a clear shift in selectivity towards mono-ARTs. The shift was structurally explained by the crystal structures, which revealed the compound induced conformational change of the catalytic residue of TNKS2. Selectivity boosts towards mono-ARTs induced by hydrophobic moieties extending towards the acceptor site has also be seen previously with PARP10 inhibitor development.^18,34^

This study is an example of a structure-based repurposing of a previously reported general inhibitor scaffold and making structure-based design of substituents enabling selectivity. In this report, we chose to study example enzymes to validate the strategy and it should be noted that there are 17 enzymes in the family and all of them have a conserved catalytic domain. Therefore, while further work is required, the strategy demonstrates ways to differentiate compound selectivity between mono- and poly-ART enzymes of the PARP family.

At the moment, very little is known about PARP15 despite that it is reported to be overexpressed in B-aggressive lymphoma^35^ and be a potential therapeutic target in acute myeloid leukemia^36^ the mechanism and contribution of this enzyme is not clear. Some of the mono-ARTs have already become validated drug targets^37^ and therefore generation of potent inhibitors with differing selectivity profiles will hopefully facilitate the studies of these less proteins and the compounds reported here may be used as such tool compounds in the future.

## Acknowledgements

The authors would like to thank the staff of the beamlines ID23-1 and ID30A-1 in ESRF and I03 in DLS. The use of the facilities of the Biocenter Oulu Structural Biology core facility, a member of Biocenter Finland, Instruct-ERIC Centre Finland and FINStruct, is gratefully acknowledged.

## Data statement

Atomic coordinates and structure factors have been deposited to the Protein Data Bank under accession numbers 7OLJ, 7OM1, 7OMC, 7OQQ, 7OSP, 7OSS, 7OSX, 7OTF, 7OTH, 7OUW and 7OUX, and raw diffraction images are available at IDA (https://doi.org/10.23729/3942aada-8d3a-4391-87ed-2a88cf624d25).

## Supplementary material

**Table S1**. Protein constructs and PARP activity assay conditions

**Table S2**. Crystallography data processing and refinement statistics

**Figure S1** ^1^H NMR spectrum of **9** in CDCl_3_ (0–15 ppm)

**Figure S2** Local zoom (6.9–7.75 ppm) of ^1^H NMR spectrum of compound **9** in CDCl_3_

**Figure S3** ^13^C NMR spectrum of **9** in CDCl_3_ (0–220 ppm)

**Figure S4** ^1^H NMR spectrum of compound **10** in CDCl_3_ (0–15 ppm)

**Figure S5** Local zooms of ^1^H NMR spectra of compound **10** in CDCl_3_ (6.9–8.6 ppm) and (CD_3_)_2_SO (6.9–8.2 ppm)

**Figure S6** ^13^C NMR spectrum of **10** in (CD_3_)_2_SO (0–220 ppm)

