## Supplementary information for "Analogs of TIQ-A as inhibitors of human mono-ADP-ribosylating PARPs"

**CONTENT**

**Table S1**. Protein constructs and PARP activity assay conditions

**Table S2**. Crystallography data processing and refinement statistics

**Figure S1** $^{1}$H NMR spectrum of **9** in $CDCl_3$ (0–15 ppm)

**Figure S2** Local zoom (6.9–7.75 ppm) of $^{1}$H NMR spectrum of compound **9** in $CDCl_3$

**Figure S3** $^{13}$C NMR spectrum of **9** in $CDCl_3$ (0–220 ppm)

**Figure S4** $^{1}$H NMR spectrum of compound **10** in $CDCl_3$ (0–15 ppm)

**Figure S5** Local zooms of $^{1}$H NMR spectra of compound **10** in $CDCl_3$ (6.9–8.6 ppm) and $(CD_3)_2SO$ (6.9–8.2 ppm)

**Figure S6** $^{13}$C NMR spectrum of **10** in $(CD_3)_2SO$ (0–220 ppm)

**Table S1**. Protein constructs and PARP activity assay conditions

| **PARP** | **Amino acids** | **Vector** | **Used in crystallization** | **Assay conditions** |
|---|---|---|---|---|
| PARP2 | 1 - 583 | pNH-TrxT | - | 30 nM PARP2 and 500 nM $NAD^+$ in a buffer consisting of 50 mM Tris pH 8.0, 5 mM $Mg^{2+}$, 10 µg/ml activated DNA, 0.1 mg/ml BSA, 0.5 h shaking at RT. |
| TNKS2 | 873 – 1161 | pNIC-MBP | - | 20 nM TNKS2 and 500 nM $NAD^+$ in a buffer consisting of 50 mM Bis-tris propane pH 7.0, 0.5 mM TCEP, 20 h shaking at RT. |
| TNKS2 | 946-1161 | pNIC-MBP | yes | - |
| PARP10 | 809 - 1017 | pNIC-CH | - | 100 nM PARP10, 2 µM SRPK2 and 500 nM Tris-HCl pH 7.0. 13 h shaking at RT. |
| PARP14 | 1535 - 1801 | pNH-TrxT | - | 500 nM PARP14 and 500 nM $NAD^+$ in a buffer consisting of 50 mM sodium phosphate pH 7, 21 h shaking at RT. |
| PARP15 | 482 - 678 | pNIC28-Bsa4 | yes | 200 nM PARP15 and 500 nM SRPK2 and 250 nM $NAD^+$ in a buffer consisting of 50 mM sodium phosphate pH 7, 3 h shaking at RT. |

**Table S2. Data collection and refinement statistics.**

| Item | PARP15-1 (PDB id. 7OQQ) | PARP15-2 (PDB id. 7OSP) | PARP15-4 (PDB id. 7OSS) | PARP15-5 (PDB id. 7OSX) |
|---|---|---|---|---|
| **Data collection** | | | | |
| Resolution (Å) | 50 – 2.00 | 50 – 1.44 | 50 – 1.50 | 50 – 1.60 |
| (Outer shell)* | (2.05 – 2.00) | (1.48 – 1.44) | (1.54 – 1.50) | (1.64 – 1.60) |
| Wavelenght | 0.97625 | 0.96600 | 0.97625 | 0.97625 |
| Beamline | ESRF, ID23-1 | ESRF, ID30A-1 | DLS, I03 | DLS, I03 |
| Temperature (K) | 100 | 100 | 100 | 100 |
| Space group | P21 21 21 | P21 21 21 | P21 21 21 | P21 21 21 |
| Cell dimensions | | | | |
| $a$, $b$, $c$ (Å) | 45.28, 68.54, 159.80 | 45.31, 68.55, 159.63 | 45.46, 68.74, 160.68 | 45.37, 68.76, 161.07 |
| α, β, γ (°) | 90, 90, 90 | 90, 90, 90 | 90, 90, 90 | 90, 90, 90 |
| No. of unique reflections | 33180 (2413) | 88158 (6625) | 81557 (5918) | 66336 (4810) |
| $R_{merge}$ | 0.095 (0.564) | 0.064 (1.120) | 0.051 (1.454) | 0.089 (1.099) |
| Mean $I/\sigma I$ | 9.05 (2.03) | 13.05 (1.45) | 25.77 (1.94) | 15.41 (2.58) |
| Completeness (%) | 96.1 (96.1) | 96.9 (100) | 100 (100) | 98.3 (97.6) |
| Redundancy | 3.57 (2.85) | 5.64 (5.70) | 13.22 (13.67) | 10.97 (10.24) |
| $CC_{1/2}$ (%) | 99.5 (74.6) | 99.9 (61.8) | 100 (74.9) | 99.8 (75.7) |
| Wilson B-factor (Å$^2$) | 23.1 | 18.9 | 21.9 | 20.8 |
| **Refinement** | | | | |
| $R_{work}$ / $R_{free}$ | 0.215 / 0.256 | 0.161 / 0.193 | 0.153 / 0.192 | 0.198 / 0.223 |
| No.all atoms | 3353 | 3571 | 3522 | 3468 |
| protein | 3234 | 3294 | 3269 | 3234 |
| ligands/ions | 22 | 30 | 26 | 19 |
| waters | 97 | 247 | 227 | 215 |
| $B$-factors (Å$^2$) | | | | |
| protein | 29.2 | 23.9 | 28.5 | 24.9 |
| ligands/ions | 40.1 | 28.5 | 31.2 | 32.4 |
| waters | 26.4 | 30.4 | 34.7 | 29.8 |
| RMSD bonds / angles | 0.009 / 1.619 | 0.010 / 1.607 | 0.008 / 1.427 | 0.008 / 1.449 |
| Ramachandran plot (%) | | | | |
| favored/allowed/outliers | 98.0 / 2.0 / 0 | 98.4 / 1.3 / 0.3 | 98.5 / 1.0 / 0.5 | 98.0 / 1.5 / 0.5 |

*Values in parentheses are for highest-resolution shell.

**Table S1. Data collection and refinement statistics. Continued**

| Item | PARP15-6 (PDB id. 7OTF) | PARP15-8 (PDB id. 7OUW) | PARP15-10 (PDB id. 7OUX) | PARP15-7 (PDB id. 7OTH) |
|---|---|---|---|---|
| **Data collection** | | | | |
| Resolution (Å) | 50 – 1.30 | 50 – 1.60 | 50 – 1.95 | 50 – 1.70 |
| (Outer shell) | (1.33 – 1.30) | (1.64 – 1.60) | (2.00 – 1.95) | (1.74 – 1.70) |
| Wavelenght | 0.97625 | 0.97625 | 0.97625 | 0.97625 |
| Beamline | DLS, I03 | DLS, I03 | DLS, I03 | DLS, I03 |
| Temperature (K) | 100 | 100 | 100 | 100 |
| Space group | P21 21 21 | P21 21 21 | P21 21 21 | P21 21 21 |
| Cell dimensions | | | | |
| *a*, *b*, *c* (Å) | 45.31, 68.63, 158.65 | 45.31, 68.89, 161.25 | 45.17, 68.58, 159.26 | 45.28, 68.70, 160.91 |
| α, β, γ (°) | 90, 90, 90 | 90, 90, 90 | 90, 90, 90 | 90, 90, 90 |
| No. of unique reflections | 122067 (8626) | 67534 (4902) | 36962 (2657) | 56195 (4118) |
| $R_{merge}$ | 0.047 (1.164) | 0.082 (1.510) | 0.142 (1.248) | 0.081 (0.824) |
| Mean $I/\sigma I$ | 23.24 (1.62) | 17.41 (1.63) | 11.71 (1.84) | 11.96 (1.22) |
| Completeness (%) | 99.7 (96.6) | 99.9 (99.1) | 100 (100) | 99.9 (99.3) |
| Redundancy | 13.01 (10.02) | 11.99 (10.19) | 13.41 (13.54) | 6.13 (3.53) |
| $CC_{1/2}$ (%) | 100 (72.0) | 99.9 (63.8) | 99.8 (81.8) | 99.9 (57.1) |
| Wilson B-factor (Å$^2$) | 18.0 | 21.5 | 29.8 | 24.6 |
| **Refinement** | | | | |
| $R_{work}$ / $R_{free}$ | 0.148 / 0.173 | 0.234 / 0.262 | 0.180 / 0.215 | 0.190 / 0.214 |
| No.all atoms | 3699 | 3314 | 3409 | 3509 |
| protein | 3330 | 3192 | 3217 | 3267 |
| ligands/ions | 27 | 20 | 21 | 22 |
| waters | 342 | 102 | 171 | 220 |
| *B*-factors (Å$^2$) | | | | |
| protein | 23.11 | 28.56 | 36.45 | 29.47 |
| ligands/ions | 22.38 | 29.21 | 38.33 | 42.19 |
| waters | 32.58 | 28.80 | 37.99 | 33.41 |
| RMSD bonds / angles | 0.010/ 1.600 | 0.007/ 1.436 | 0.007/ 1.411 | 0.007/ 1.619 |
| Ramachandran plot (%) favored/allowed/outliers | 98.2 / 1.6 / 0.2 | 97.5 / 2.2 / 0.3 | 97.5 / 2.0 / 0.5 | 97.5 / 2.2 / 0.3 |

*Values in parentheses are for highest-resolution shell.

**Table S2. Data collection and refinement statistics. Continued**

| Item | TNKS2-8 (PDB 7OM1) | TNKS2-10 (PDB 7OMC) | TNKS2-7 (PDB 7OLJ) |
|---|---|---|---|
| **Data collection** | | | |
| Resolution (Å) | 45.39 - 1.70 | 45.35 - 2.10 | 42.49 - 1.80 |
| (Outer shell) | 1.761 - 1.70 | 2.175 - 2.10 | 1.864 - 1.80 |
| Wavelenght | 0.97625 | 0.97625 | 0.97625 |
| Beamline | DLS, I03 | DLS, I03 | DLS, I03 |
| Temperature (K) | 100 | 100 | 100 |
| Space group | C222(1) | C222(1) | C222(1) |
| Cell dimensions | | | |
| *a*, *b*, *c* (Å) | 91.21, 98.3, 118.27 | 90.69, 98.18, 117.48 | 90.93 98.53 119.37 |
| α, β, γ (°) | 90, 90, 90 | 90, 90, 90 | 90, 90, 90 |
| No. of unique reflections | 58597 (5791) | 30936 (3040) | 49857 (4954) |
| $R_{merge}$ | 0.05454 (0.7849) | 0.074 (1.189) | 0.06333 (0.9564) |
| Mean $I/\sigma I$ | 19.01 (2.09) | 22.06 (2.29) | 17.28 (1.76) |
| Completeness (%) | 99.89 (99.97) | 99.95 (99.93) | 99.82 (99.96) |
| Redundancy | 6.7 (6.4) | 13.5 (14.0) | 6.8 (6.6) |
| $CC_{1/2}$ (%) | 0.999 (0.817) | 0.999 (0.814) | 0.999 (0.759) |
| Wilson B-factor (Å$^2$) | 27.54 | 47.16 | 32.60 |
| **Refinement** | | | |
| $R_{work}$ / $R_{free}$ | 0.1854 / 0.1981 | 0.184 / 0.219 | 0.1915 / 0.2113 |
| No.all atoms | 3811 | 3622 | 3728 |
| protein | 3388 | 3349 | 3376 |
| ligands/ions | 66 | 62 | 64 |
| waters | 357 | 211 | 288 |
| *B*-factors (Å$^2$) | | | |
| protein | 32.77 | 53.70 | 36.74 |
| ligands/ions | 36.37 | 59.97 | 39.25 |
| waters | 39.54 | 55.89 | 40.91 |
| RMSD bonds / angles | 0.013 / 1.61 | 0.013 / 1.65 | 0.013 / 1.63 |
| Ramachandran plot (%) | | | |
| favored/allowed/outliers | 98.5 / 1.5 / 0 | 98.3 / 1.7 / 0 | 99.0 / 1.0 / 0 |

**Figure S1** $^{1}$H NMR spectrum of compound **9** in $CDCl_3$ (0–15 ppm)

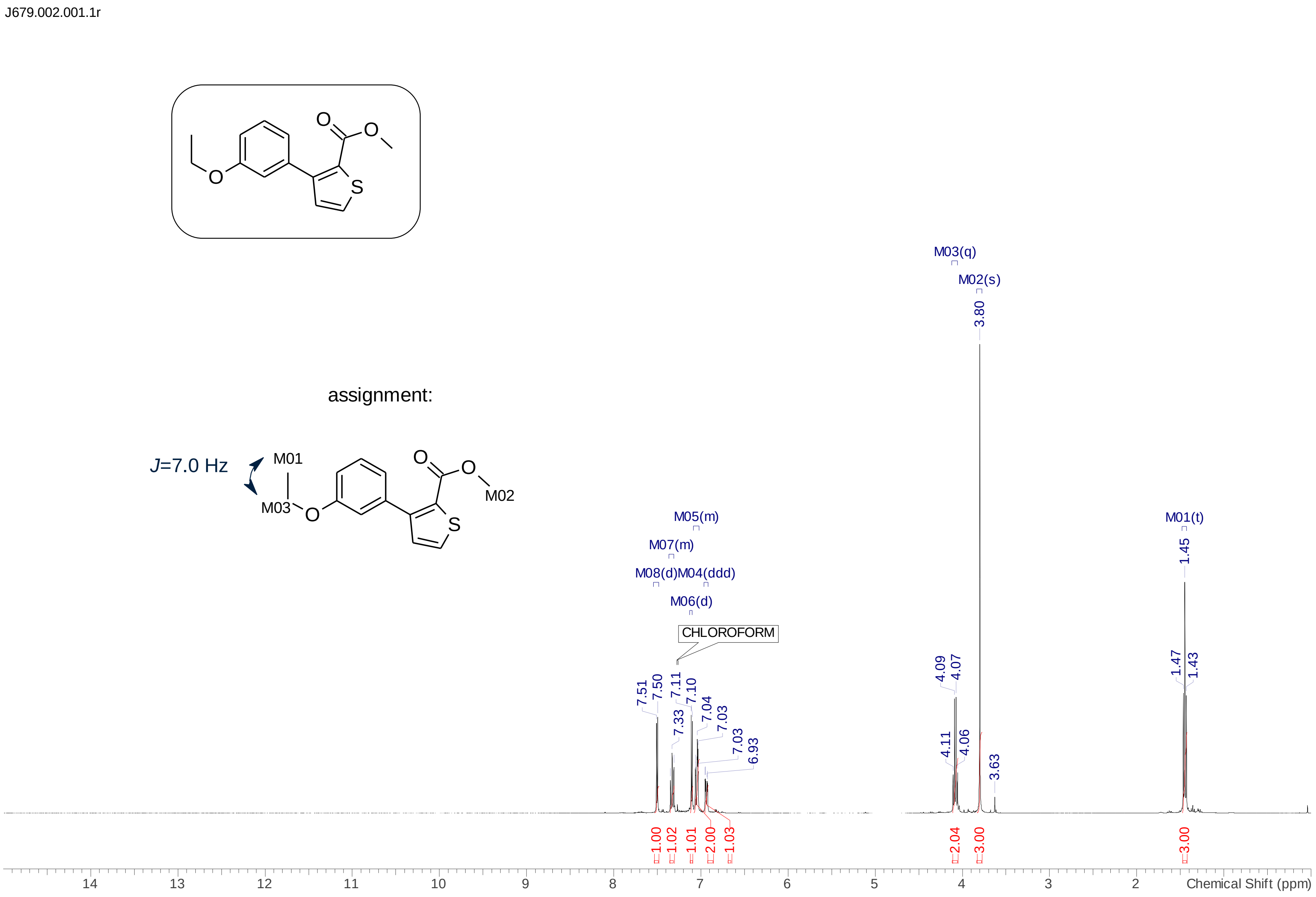

**Figure S2** Local zoom (6.9–7.75 ppm) of [1]H NMR spectrum of compound **9** in $CDCl_3$

J679.002.001.1r

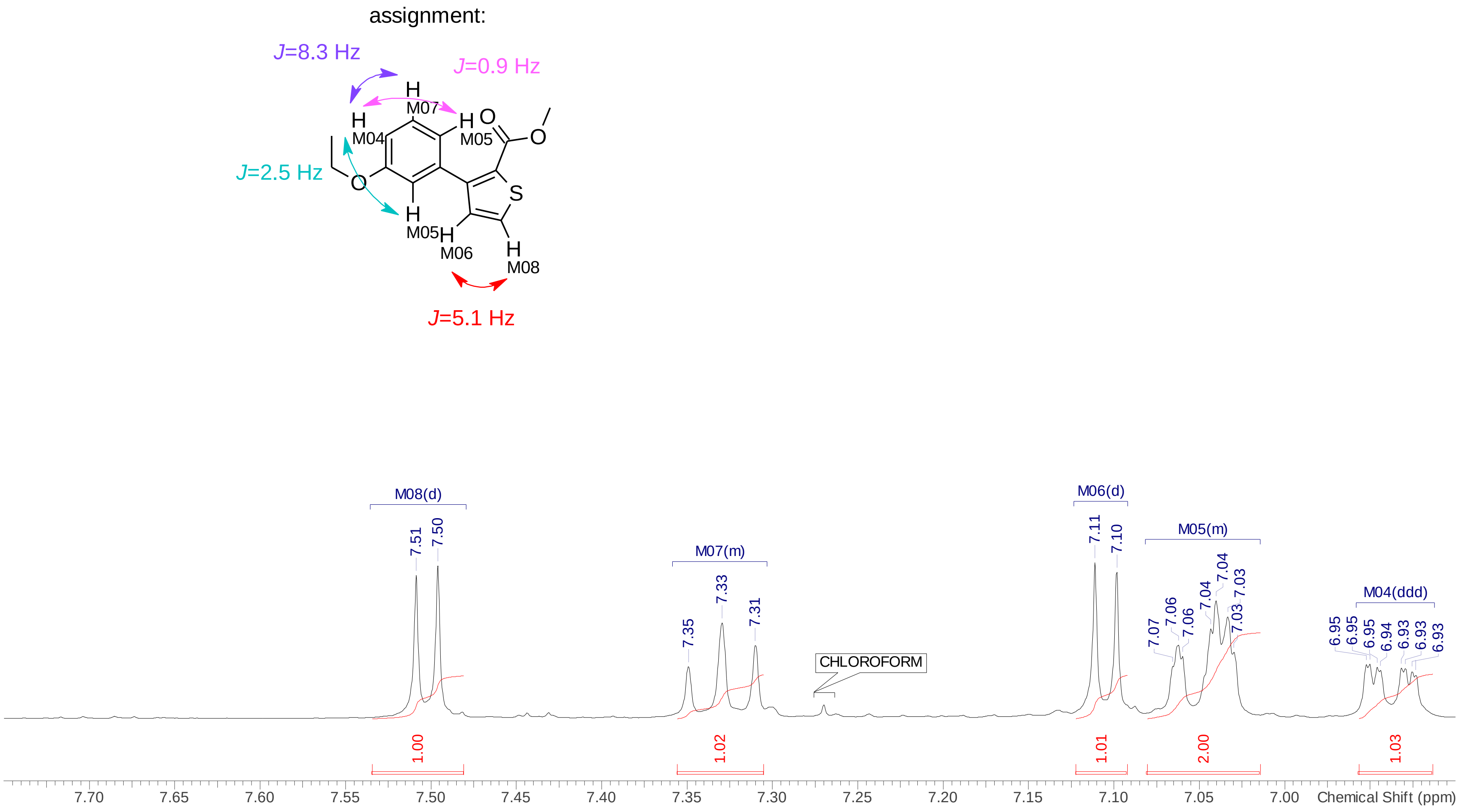

**Figure S3** $^{13}$C NMR spectrum of **2** in $CDCl_3$ (0–220 ppm)

J679.003.001.1r

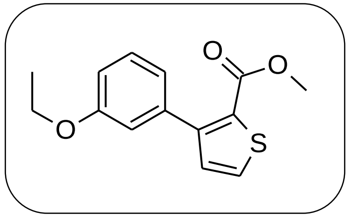

| No. | (ppm) | (Hz) | Height |
|---|---|---|---|
| 1 | 14.75 | 1484.5 | 0.7595 |
| 2 | 51.79 | 5211.1 | 0.3190 |
| 3 | 63.29 | 6368.0 | 0.5694 |
| 4 | 113.97 | 11467.2 | 0.8449 |
| 5 | 115.32 | 11602.2 | 0.7996 |
| 6 | 121.48 | 12222.1 | 0.8303 |
| 7 | 126.86 | 12763.5 | 0.2311 |
| 8 | 128.67 | 12945.4 | 1.0000 |
| 9 | 130.07 | 13086.3 | 0.9219 |
| 10 | 131.45 | 13225.7 | 0.8776 |
| 11 | 136.78 | 13761.9 | 0.5541 |
| 12 | 148.34 | 14924.7 | 0.4607 |
| 13 | 158.32 | 15929.0 | 0.4972 |
| 14 | 162.26 | 16325.1 | 0.3340 |

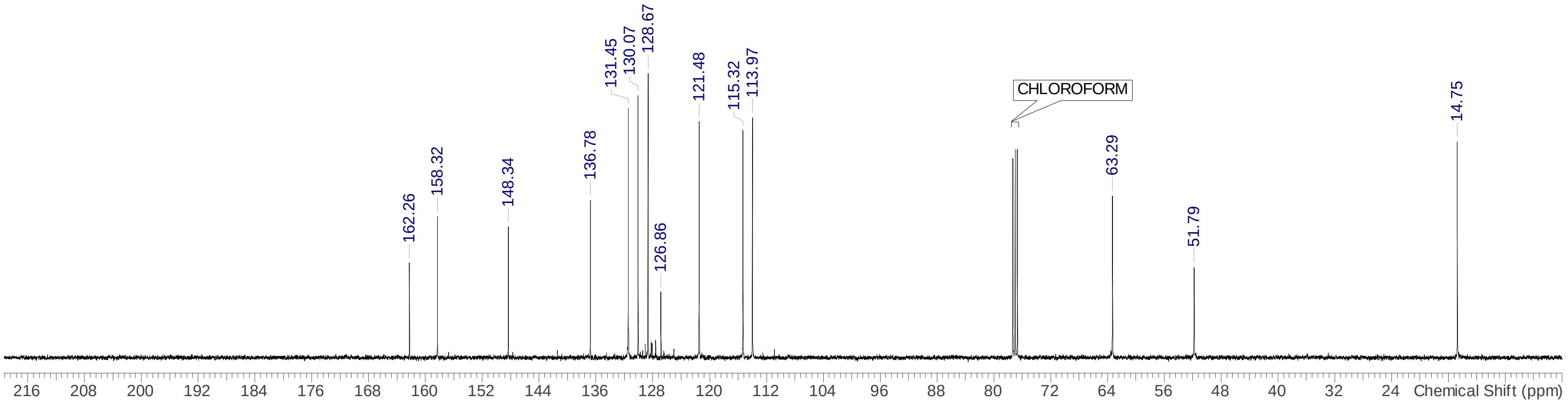

**Figure S4** $^{1}$H NMR spectrum of compound **10** in $CDCl_3$ (0–15 ppm)

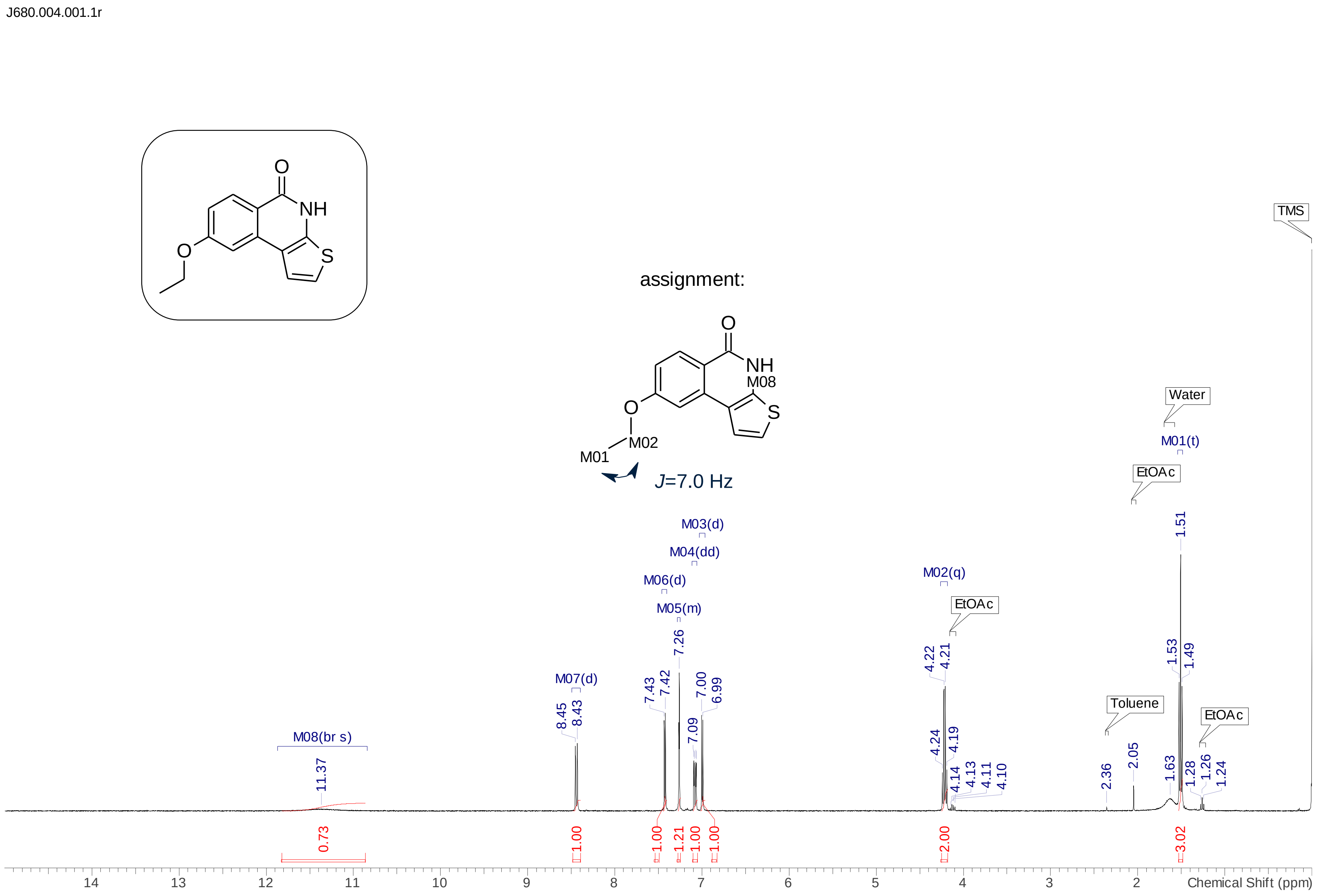

**Figure S5** Local zooms of $^{1}$H NMR spectra of compound **10** in $CDCl_3$ (the upper; 6.9–8.6 ppm) and $(CD_3)_2SO$ (the lower; 6.9–8.2 ppm)

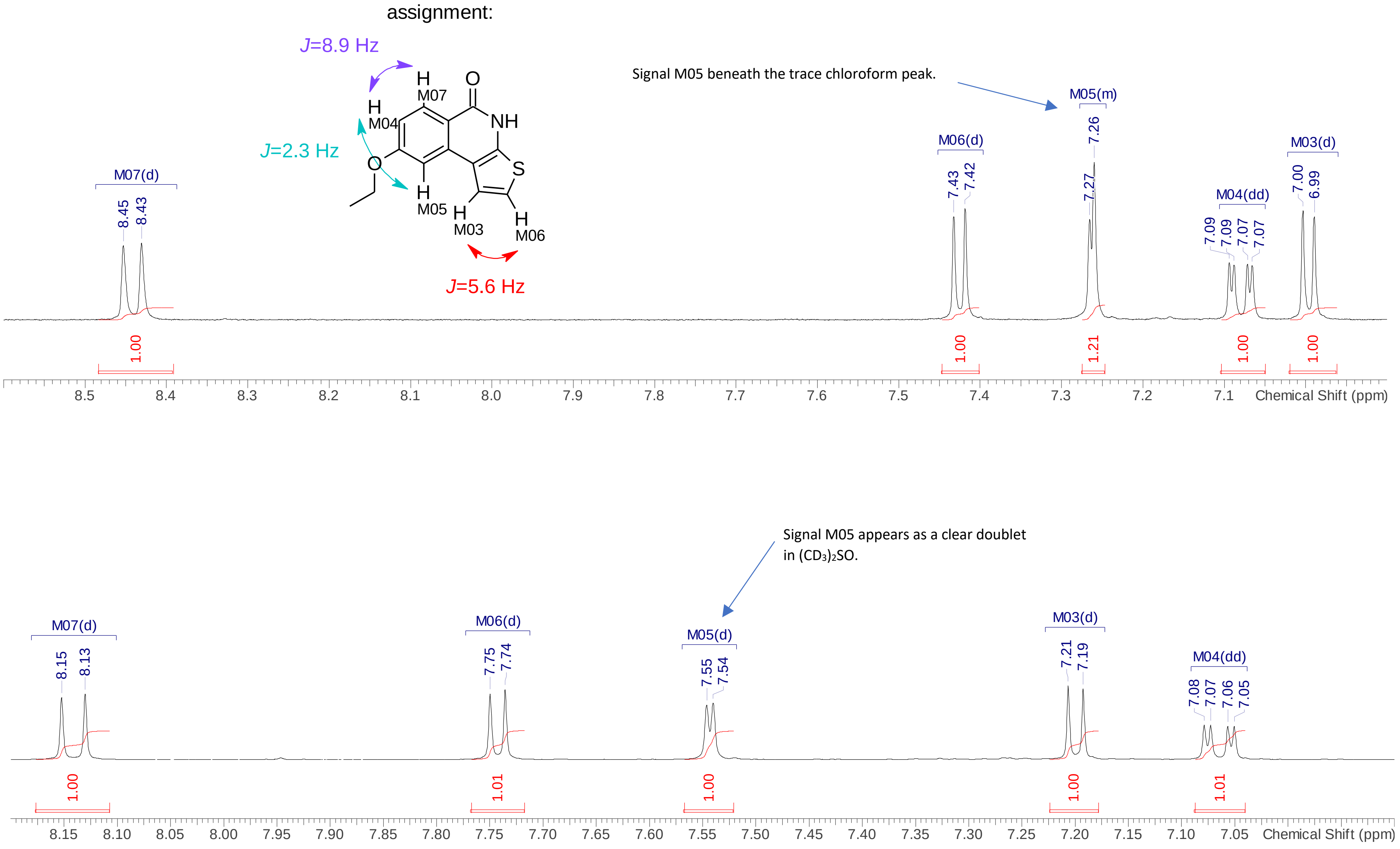

**Figure S6** $^{13}C$ NMR spectrum of **10** in $(CD_3)_2SO$ (0–220 ppm)

J680.003.001.1r

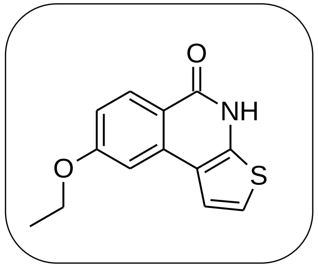


| No. | (ppm) | (Hz) | Height | No. | (ppm) | (Hz) | Height | No. | (ppm) | (Hz) | Height | No. | (ppm) | (Hz) | Height |
|---|---|---|---|---|---|---|---|---|---|---|---|---|---|---|---|
| 1 | 14.54 | 1462.7 | 0.0410 | 5 | 116.24 | 11695.6 | 0.0319 | 9 | 129.94 | 13073.3 | 0.0370 | 13 | 162.17 | 16316.5 | 0.0297 |
| 2 | 63.71 | 6410.0 | 0.0273 | 6 | 116.86 | 11757.9 | 0.0229 | 10 | 135.61 | 13644.0 | 0.0251 | | | | |
| 3 | 105.64 | 10628.9 | 0.0401 | 7 | 117.62 | 11834.2 | 0.0285 | 11 | 141.31 | 14217.7 | 0.0129 | | | | |
| 4 | 114.97 | 11567.9 | 0.0358 | 8 | 121.52 | 12226.7 | 0.0425 | 12 | 160.98 | 16196.9 | 0.0368 | | | | |

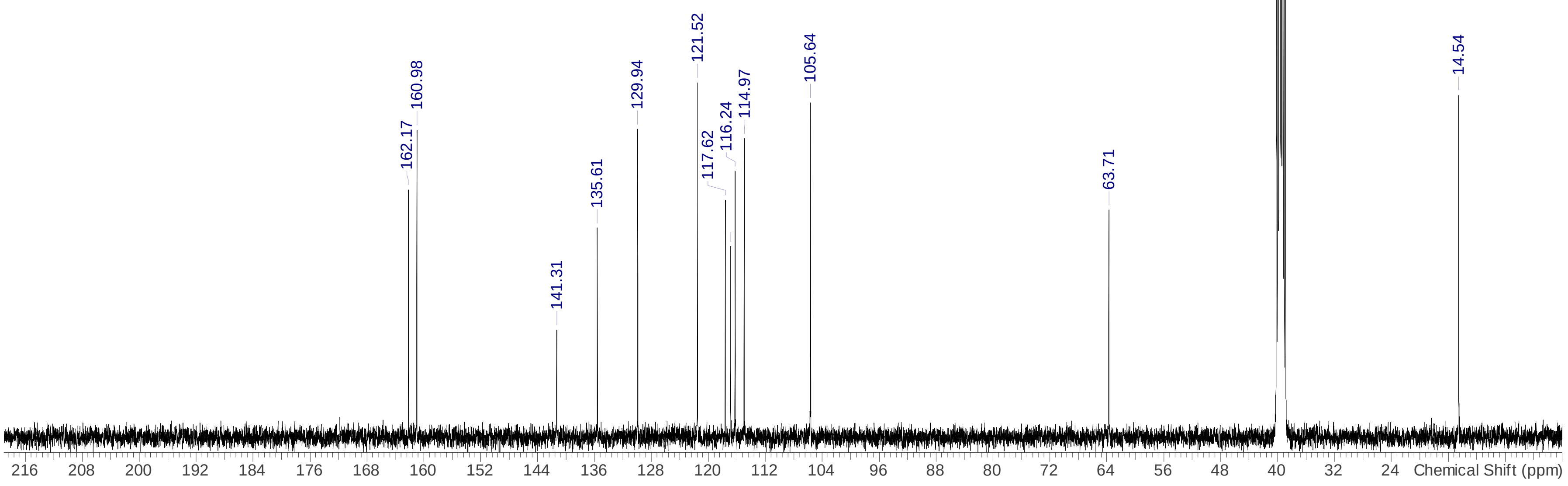
